# The frequency of cortical microstimulation shapes artificial touch

**DOI:** 10.1101/659516

**Authors:** Thierri Callier, Nathan Brantly, Attilio Caravelli, Sliman J. Bensmaia

## Abstract

Intracortical microstimulation (ICMS) of somatosensory cortex evokes vivid tactile sensations and can be used to convey sensory feedback in brain-controlled bionic hands. Changes in ICMS frequency result in discriminable percepts, but the effects of frequency have only been investigated over a narrow range of low frequencies, spanning only a small fraction of that relevant for neuroprosthetics. Furthermore, the sensory correlates of changes in ICMS frequency remain to be elucidated. To fill these gaps, we trained monkeys to discriminate the frequency of ICMS pulse trains over a wide range of frequencies (from 10 to 400 Hz). ICMS amplitude also varied across stimuli to reduce the animals’ reliance on magnitude in making frequency judgments. We found that animals could discriminate ICMS frequency up to about 200 Hz but that the sensory correlates of frequency were highly electrode dependent. We discuss the implications of our findings for neural coding and brain-machine interfaces.

## Introduction

Intracortical microstimulation (ICMS) in somatosensory cortex has been shown to evoke tactile percepts the location and magnitude of which can be systematically manipulated (Berg et al., 2013; Flesher et al., 2016; Kim et al., 2015; Tabot et al., 2013). This phenomenon can be exploited to convey sensory feedback in sensorized bionic hands: the output of sensors on the prosthetic fingers can drive stimulation through electrodes located in the appropriate region of the somatosensory homunculus – thereby intuitively conveying information about the location on the hand of contact with objects – and the strength of stimulation can be modulated to produce sensations whose magnitude depends on the output of the sensor – thereby intuitively conveying information about pressure at each contact location (Tabot et al., 2013). Ideally, the electrically induced neuronal activity would mimic its mechanically induced counterpart in able bodied individuals, which would lead to completely natural sensation (Bensmaia, 2015). However, limitations inherent to electrical stimulation, in the number of stimulating channels, and in our understanding of cortical circuitry severely restrict our ability to produce naturalistic neuronal activity.

Despite its unnaturalness, however, ICMS leads to sensations that are reported by human subjects as being natural or nearly so (Flesher et al., 2016; Salas et al., 2018) and ICMS-based feedback leads to improved functionality for brain-controlled bionic hands (Romo et al., 2000, 1998). In light of this, we seek to refine our understanding of how the parameters of ICMS shape the percept to produce increasingly intuitive and useful artificial touch. As alluded to above, the effects of ICMS amplitude on the evoked sensations have been extensively studied, as has the effect of stimulation location on the cortical sheet (i.e., through different electrodes) (Berg et al., 2013; Flesher et al., 2016; Kim et al., 2015; Salas et al., 2018; Tabot et al., 2013). Changes in ICMS frequency have been shown to evoke discriminable percepts in studies with non-human primates (Romo et al., 2000, 1998). However, the range of frequencies tested only spanned a small fraction of that relevant for neuroprosthetics (from 10 to 30 Hz).

Furthermore, the question remains *how* changes in frequency affect the evoked percept. Indeed, sensitivity to ICMS increases with frequency (Kim et al., 2015), so the animals’ ability to discriminate frequency may rely on frequency-dependent changes in perceived magnitude. Alternatively, changes in frequency may lead to differences in the quality of the percept. Of course, these two possibilities are not mutually exclusive.

The objective of the present study, then, was to characterize the ability of non-human primates (Rhesus macaques) to discriminate ICMS applied to somatosensory cortex over a wide range of frequencies (from 10 to 400 Hz) and to assess the degree to which changes in frequency affect the magnitude of the percept, its quality, or both.

## Results

We implanted three monkeys with electrode arrays (Utah electrode arrays, UEAs) in Brodmann’s area 1 of somatosensory cortex and had them judge which of two 1-s long ICMS pulse trains was higher in pulse frequency (Figure 1). On each experimental block, consisting of several hundred trials, a standard stimulus (at 20, 50, 100, or 200 Hz) was paired with several comparison stimuli whose frequencies varied around the standard frequency. The amplitudes of the standard and comparison stimuli also varied from trial to trial (50, 60, 70 or 80 µA presented in every possible combination in random order). The animal was rewarded when it reported which of the two stimuli was higher in frequency. The large, behaviorally irrelevant variations in amplitude were intended to reduce or abolish the informativeness of perceived magnitude, which is modulated by changes in both frequency and amplitude.

**Figure 1:**
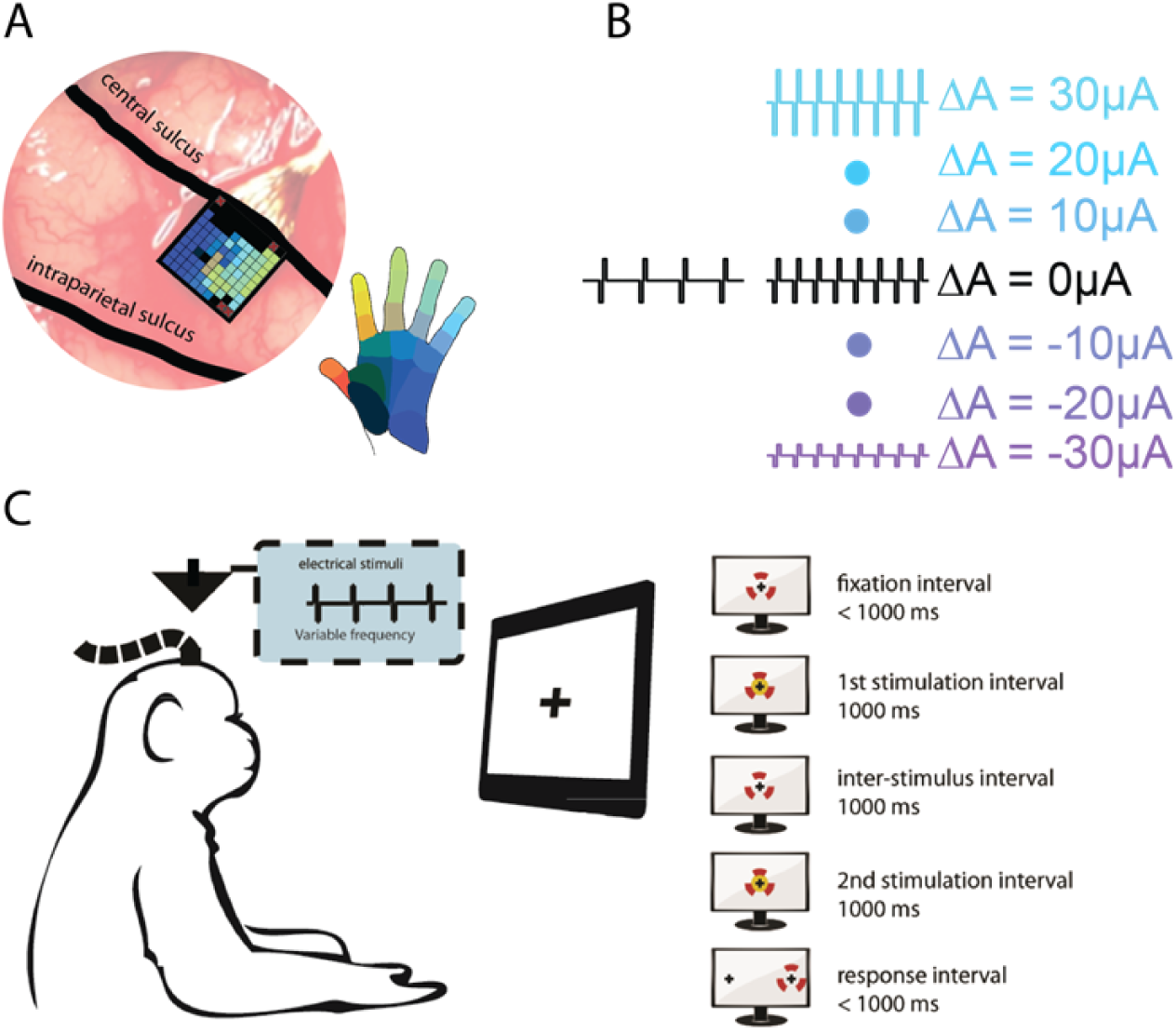
Experimental design. A| A Utah Electrode Array (UEA) was implanted in the hand representation of area 1 (Monkey C). B| The animal’s task was to select the pulse train with the higher frequency and ignore changes in stimulus amplitude. C | Left: The animal was seated facing a monitor while ICMS was delivered through the UEA. Right: trial sequence. The red markers denote the gaze of the animal, which fixated on a target in the center of the screen while a pair of ICMS pulse trains was sequentially delivered, then reported its judgment by saccading to one of two targets.

### Frequency discrimination with equal amplitudes

On all electrodes tested, the animals were able to reliably discriminate the frequency of ICMS pulse trains when the amplitudes of the standard and comparison were equal, except over the highest frequency range (from 200 to 400 Hz)(Figure 2A). Indeed, the animals reached near perfect performance with the 20-, 50-, and 100-Hz standards and for frequencies below 200 Hz with the 200-Hz standard. However, when both frequencies were 200 Hz and above, performance leveled off, often below 75%, suggesting that further increases in frequency had no impact on the evoked sensations. To gauge the animals’ sensitivity to changes in frequency, we computed the frequency increment or decrement required to achieve 75% discrimination performance, known as the just noticeable difference (JND). We found that JNDs increased with standard frequency from around 3 Hz for a 20-Hz standard to 95-Hz for the 200 Hz standard (Figure 2B). Weber fractions – the ratio of the JND to the standard frequency – increased dramatically, from 0.15 to 0.5, between 50 and 100Hz, indicating a much higher sensitivity to changes in frequency in the low range (Figure 2C). Frequency discrimination performance was independent of stimulus amplitude at the low frequencies but improved with increasing amplitude at the high frequencies (Figure 2A, Supplementary figure 1). Note that, while only trials with equal amplitudes were included in these results, these trials were interleaved with many more trials on which the standard and comparison amplitudes differed. The monkeys’ strategy was therefore geared towards selecting the higher frequency in the face of intensive confounds. If trained with only equal amplitude pairs, animals may have exploited intensity differences and performed better.

**Figure 2:**
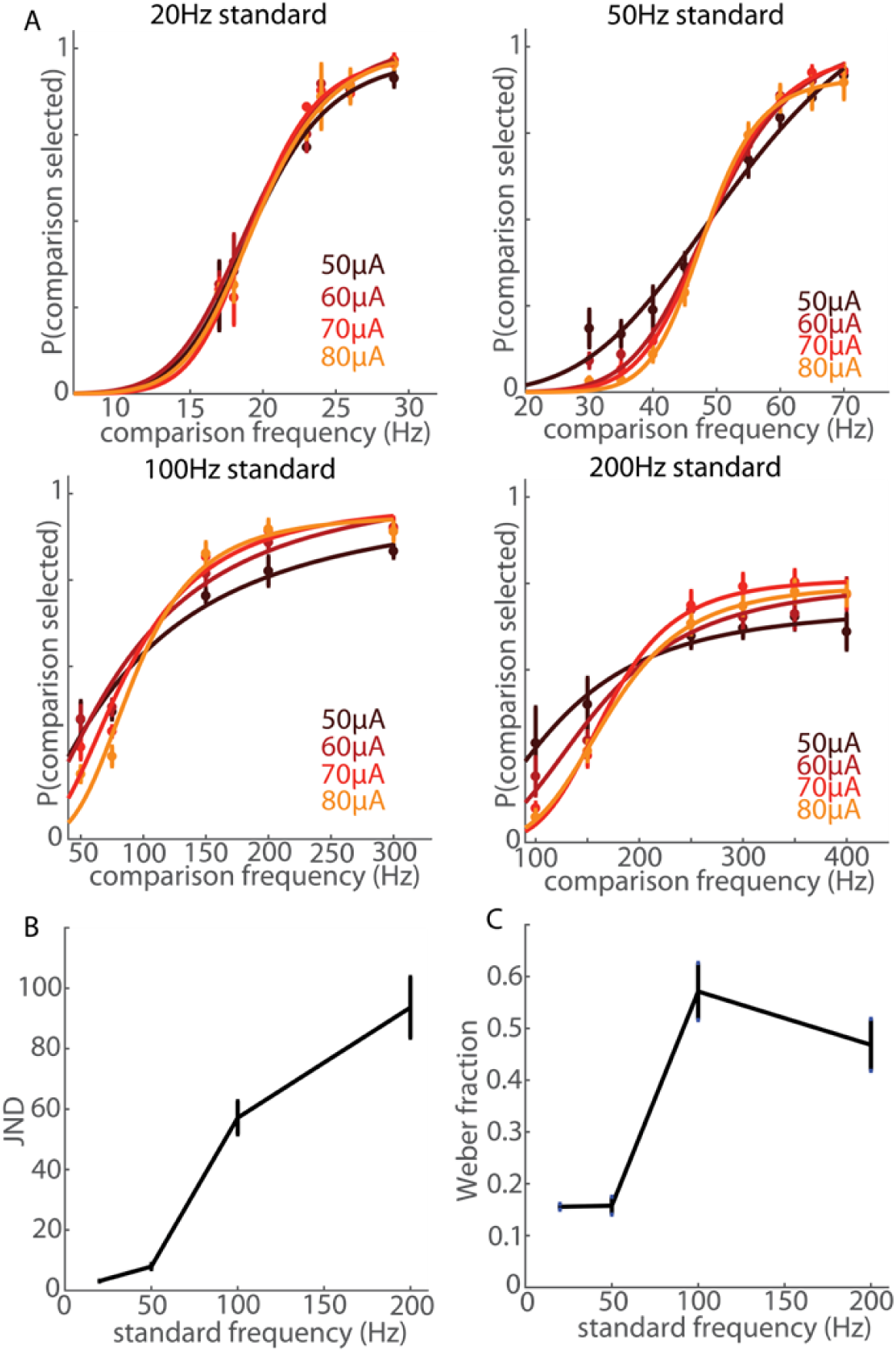
Frequency discrimination with equal amplitudes. A| Performance on the frequency discrimination task when stimulus amplitudes were equal, averaged across all electrodes (N = 1 each from monkeys A and B for the 20-Hz standard, N = 5 from monkeys A and B for the 50-Hz and 200Hz standard, N = 8 from monkeys A, B, and C for the 100-Hz standard Different colors denote different stimulus amplitudes. The animals achieved high performance for frequencies below 200 Hz. B| Just-noticeable-difference (JND) as a function of standard frequency. JNDs at each amplitude were averaged for each electrode. Error bars denote the standard error of the mean across the all electrodes tested at each standard. C| Weber fractions as a function of standard frequency.

### Frequency discrimination with unequal amplitudes

One central objective of the present study was to assess the extent to which changes in frequency shape the evoked percept beyond modulating its magnitude. Indeed, one might expect higher microstimulation frequencies to evoke stronger sensations given the increased sensitivity at higher frequencies as reflected in the detection thresholds (Kim et al., 2015). However, increases in microstimulation amplitude (charge per pulse) also evoke stronger sensations (Flesher et al., 2016; Tabot et al., 2013). To the extent that frequency discrimination judgments were based on intensive differences, then, we expected the psychometric functions to shift to lower or higher frequencies (i.e. left- or rightward) depending on the amplitude difference between the standard and comparison stimulus. As expected, animals exhibited a systematic bias towards selecting the higher-amplitude stimulus (Figure 3), as evidenced by a left- or rightward shift in the psychometric functions when the standard stimulus was lower or higher in amplitude than the comparison stimulus, respectively. The direction of the bias was consistent across electrodes and frequencies and the magnitude of the bias increased monotonically as the difference in amplitude between the stimuli increased. The magnitude of the bias also varied widely across electrodes (Figure 3): on some electrodes, amplitude differences slightly contaminated frequency judgments (top row); on others, amplitude differences dominated the animal’s choices (bottom row).

**Figure 3:**
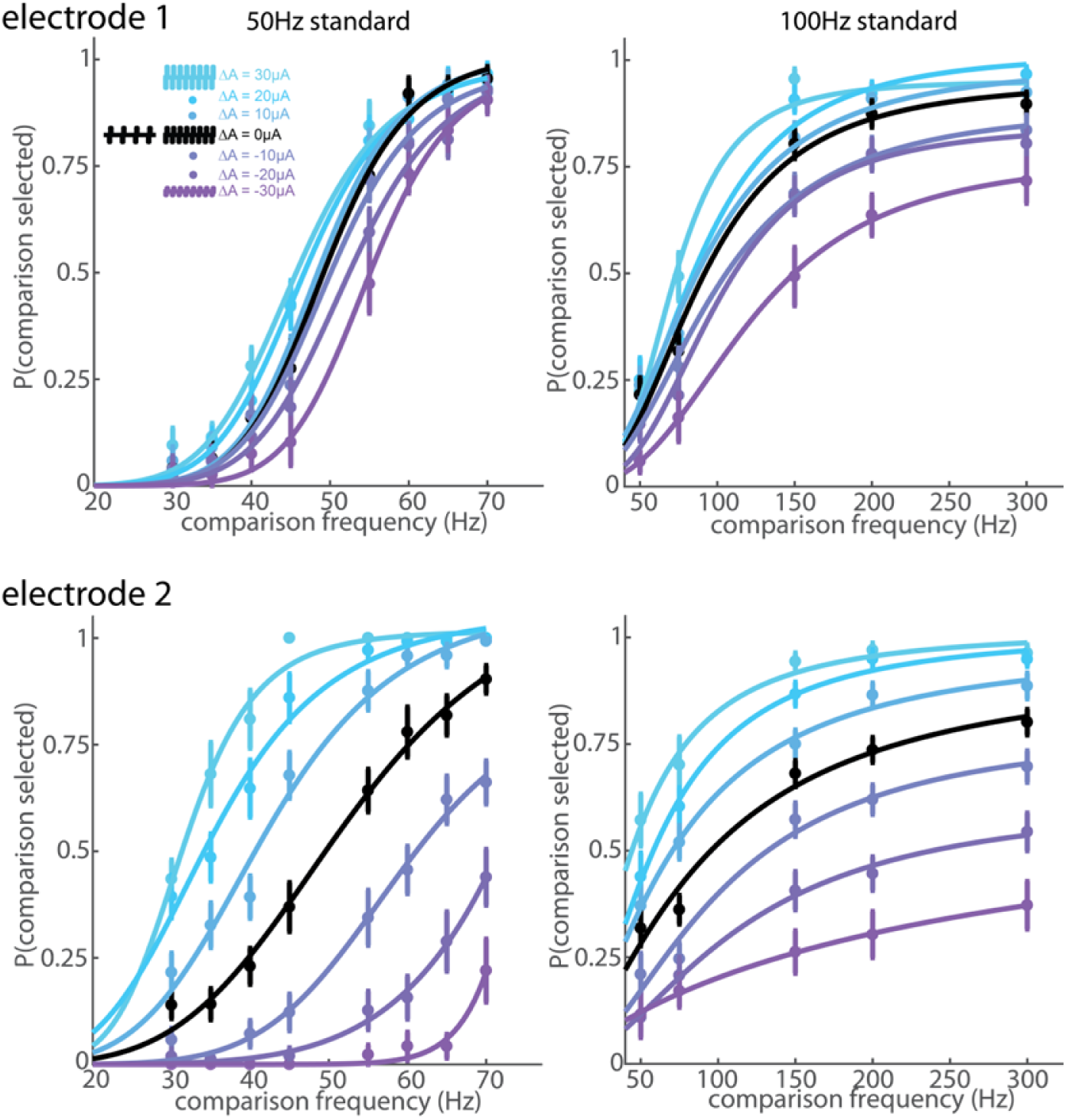
Frequency discrimination with variable amplitudes. Top row: behavioral performance at one electrode from monkey A for standard frequencies of 50 and 100Hz. Colors indicate amplitude differences between the comparison and standard frequencies (blue indicates the comparison stimulus amplitude was higher). The animal’s choices were slightly biased toward the higher amplitude. Bottom row: The same monkey’s performance for a different electrode. The monkey could perform frequency discrimination with equal amplitudes at both electrodes (black), but amplitude exerted a powerful influence on its frequency judgments when stimulation was delivered through Electrode 2.

The bias can be quantified by computing the point of subjective equality (PSE) – at which the animal is equally likely to choose the standard or comparison stimulus – for each condition. Systematic deviations of the PSE away from the standard indicate a bias. To quantify the magnitude of the bias at each electrode and standard, we computed the slope of the function relating the natural log of the PSE to amplitude difference (Supplementary figure 2). Slopes tended to be higher for higher standard frequencies but varied widely across electrodes, especially for the 50 and 100Hz standards (Supplementary figure 2B), reflecting differences in susceptibility of frequency discrimination to differences in amplitude across electrodes.

### Differences across electrodes

After extensively testing a few electrodes as described above, we had the animals perform the frequency discrimination task using a more restricted set of stimuli to sample electrodes more widely. In this set, the frequency difference between the two trial stimuli was always 100 Hz, with the low frequency stimulus spanning the range from 70 to 170 Hz, and the amplitudes were the same as those tested in the full set. The 17 electrodes tested yielded a wide range of biases (Figure 4, Supplementary figure 3): performance was consistently high when the higher-frequency stimulus was also higher in amplitude, whereas performance in the converse conditions – when the higher-frequency stimulus was lower in amplitude – varied widely. Poor performance when the high-frequency stimulus was lower in amplitude indicated a reliance on intensity differences to perform the frequency discrimination task.

**Figure 4:**
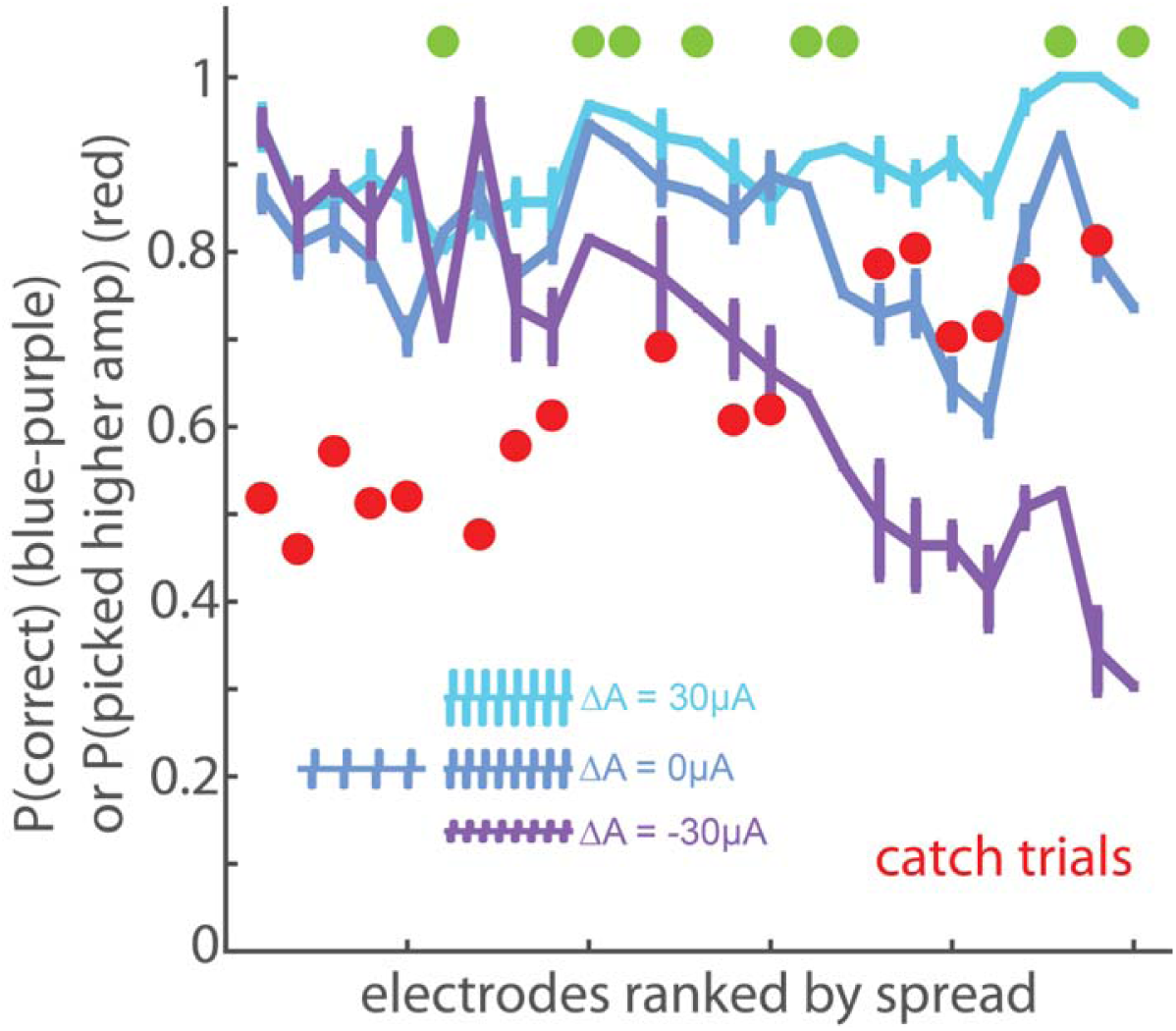
Magnitude of the amplitude bias across electrodes. For 25 tested electrodes, asymptotic performance on the frequency discrimination task with a frequency difference of 100 Hz (with lower frequency ranging from 70 to 170 Hz) and amplitude differences of −30, 0, and 30 µA. Electrodes are ranked by the difference in performance between the two amplitude extremes (cyan and purple). Green dots indicate the 100 vs 200 Hz performance for the 8 electrodes that were extensively tested (full psychometric curves were obtained). Red dots indicate the animal’s tendency to select the higher amplitude stimulus on catch trials (on which both stimuli are of equal frequency).

Next, we assessed what might account for these differences in frequency sensitivity across electrodes. ICMS frequency discrimination has been previously hypothesized to be dependent on the stimulated population: Electrodes that impinge upon cortical neurons with rapidly adapting responses (RA-like) yielded better performance than did those that impinged upon neurons with slowly adapting responses (SA-like)(Romo et al., 2000, 1998). Note that the distinction between RA- and SA-like responses has been called into question, as most cortical neurons exhibit intermediate responses (Pei et al., 2009; Saal & Bensmaia, 2014). Nonetheless, we tested this hypothesis by examining the responses to mechanical indentations delivered to the receptive field of the neurons surrounding each electrode tested. From these responses, we computed an adaptation index to gauge how “RA-like” the response was at each electrode (Callier, Suresh, and Bensmaia, 2018; Pei et al., 2009). We found no consistent relationship between RA index and the susceptibility to the amplitude confound (Supplementary figure 4).

Another possibility is that differences in discrimination performance reflect differences in sensitivity to ICMS. To test this hypothesis, we measured the detection thresholds on a subset of electrodes used in the frequency discrimination experiment and found no relationship between frequency discrimination performance and detection threshold (measured at 100 Hz, Supplementary figure 5). In other words, the poor performance on frequency discrimination on some electrodes cannot be attributed to an inability to feel the stimulation.

### Disentangling frequency and amplitude effects

Next, we assessed whether behavioral performance could be explained by a model in which the animals discriminated frequency on the basis of differences in sensory magnitude, itself driven by both microstimulation frequency and amplitude. Specifically, from amplitude discrimination experiments (with frequency held constant), we could estimate sensitivity to changes in amplitude; from frequency discrimination experiments (with amplitude held constant), we could estimate sensitivity to changes in frequency; from experiments in which both frequency and amplitude vary, we could estimate the degree to which performance was explained by a linear combination of frequency and amplitude differences (under the assumption they can be collapsed to a single sensory dimension)(Supplementary figure 6). We could then assess the degree to which the relative sensitivities to frequency and amplitude derived from the single-parameter experiments matched those derived from the combined parameter experiments. To this end, we built iso-performance contours built from behavioral data where only amplitude or only frequency varied and compared those to contours derived from behavioral data when both of these parameters varied (Supplementary figure 7, Supplementary figure 8).

The contours derived from combined variable experiments were consistently shallower than were those derived from the single-variable discrimination experiments (Supplementary figure 8). That is, the effect of frequency relative to that of amplitude was strongly underestimated from single-variable experiments, especially for electrodes with a weak amplitude bias (Supplementary figure 7). In other words, frequency differences cannot be reversed by amplitude differences of the opposite sign as readily as would be predicted from the results of single-variable experiments. These results are consistent with the hypothesis that changes in ICMS frequency not only modulate the magnitude of the sensation but also its quality.

To further test the hypothesis that frequency changes have non-intensive effects on perception, we introduced catch trials in the frequency discrimination experiment in which the two stimuli in the pair were at the same frequency but different amplitudes. To the extent that the animal relied on intensive cues to make its judgment, it would select the higher amplitude stimulus. To the extent that it judged the stimuli along a frequency-specific continuum and ignored differences in intensity (as it was rewarded to do), it would be equally likely to select either stimulus. We found that, for electrodes with a weak amplitude bias, the animal was equally likely to pick either stimulus on catch trials. For electrode with a strong amplitude bias, on which we hypothesized the animal was performing an intensity discrimination task, it was highly likely to pick the higher amplitude stimulus on catch trials (Figure 4, Supplementary figure 10, also see “validation of catch trials” in the supplementary text).

Results from simulations suggest that the perceived magnitude of a stimulus can be predicted from the population spike count (Kim et al., 2017). To the extent that animals were not solely relying on intensity differences, however, this neural code is unlikely to exclusively mediate their behavioral performance. One possibility is that the stimulus can be decoded from the spatial layout of the cortical response. For example, frequency and amplitude may each contribute systematically to the falloff in the response with distance from the electrode tip. Their individual contributions could then be untangled by sampling total evoked activity at several distances from the electrode tip. To investigate this possibility, we delivered pulse trains in which the pulse amplitude varied from pulse to pulse over a range but were on average equal to the amplitudes of constant-amplitude pulse trains. The spatial extent of the response – the pattern of recruitment – thus varied from pulse to pulse, blurring the formerly sharp separation between distinct spatial patterns of recruitment, and so reliance on a spatial pattern of activation would be lead to lower frequency discrimination performance. In individual experimental blocks, we randomly interleaved trials on which 1) both pulse trains had constant amplitudes as in previous experiments, 2) both pulse trains had variable amplitudes, and 3) one pulse train was constant in amplitude and the other was variable. The (mean or constant) amplitude of each stimulus was either the same or different as in constant-amplitude blocks. The pulse-by-pulse variability in amplitude had a negligible effect on performance (Figure 5, Supplementary figure 9), suggesting that frequency discrimination does not rely on differences in spatial patterns of electrically evoked neural activation.

**Figure 5:**
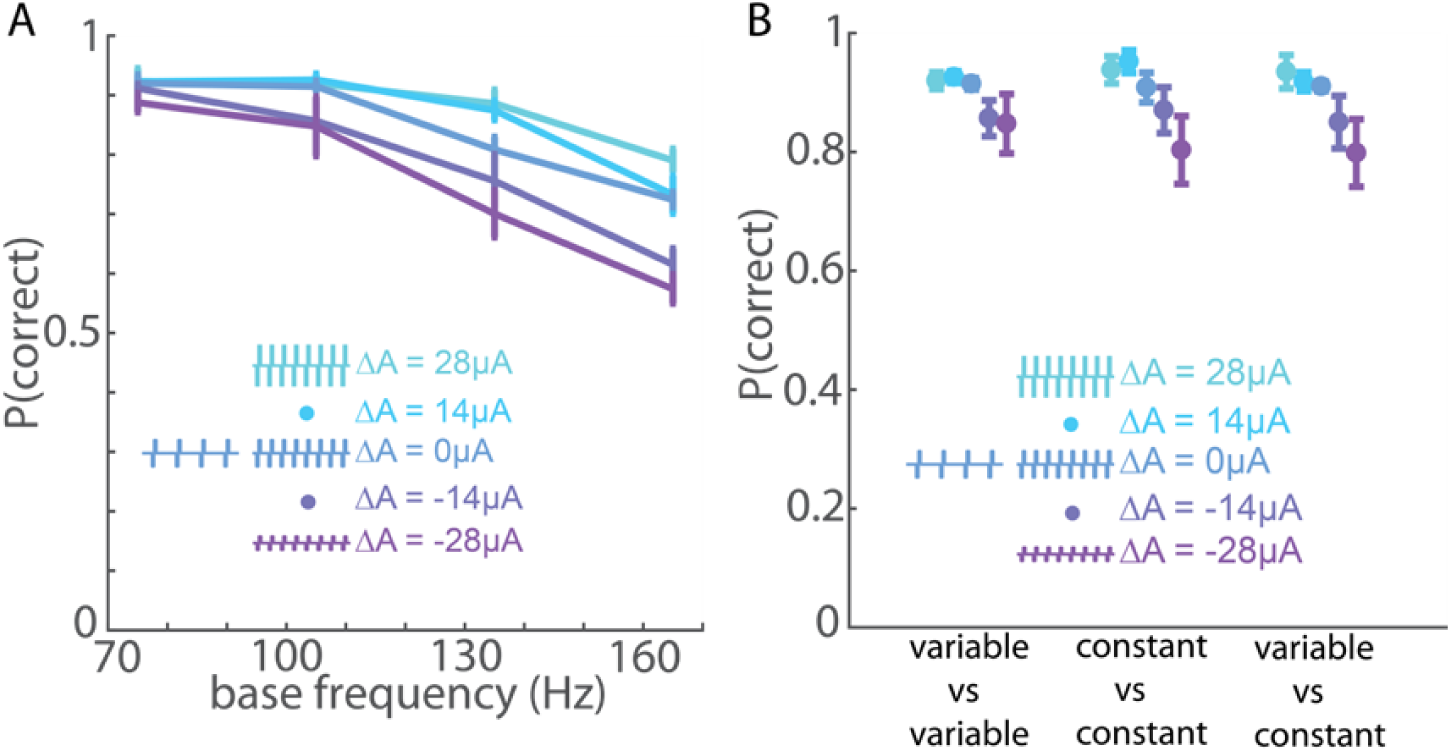
Varying individual pulse amplitude has a negligible effect on frequency discrimination performance. A| Monkey B’s performance vs. base frequency in the variable-amplitude experiment for a group of 4 electrodes with weak amplitude bias. The frequency difference was always 90 Hz for the variable-amplitude experiment due to hardware constraints (see methods). B| Performance when stimulus pulse trains were both variable-amplitude, were split, or were both constant-amplitude, for the same 4 electrodes with weak amplitude bias. Changing spatial distribution of the ICMS-induced activity on a pulse by pulse basis had little to no effect on the animal’s ability to discriminate frequency.

## Discussion

In summary, we find that (1) animals can discriminate frequency of ICMS up to about 200 Hz, (2) changes in frequency affect both the magnitude and quality of the evoked sensation, (3) the degree to which frequency shapes sensory quality varies across electrodes, and (4) ICMS frequency does not shape the percept by shaping the spatial pattern of neural activation.

### Microstimulation frequency can be discriminated up to approximately 200 Hz

When stimuli differing in frequency but matched in amplitude were paired, animals were able to discriminate frequency up to approximately 200 Hz, beyond which performance declined considerably (Figure 2, Supplementary figure 1), suggesting that increases in frequency beyond 200 Hz only weakly affect the evoked percept. This frequency cut-off coincides with the point at which detection thresholds for ICMS level off (Kim et al., 2015).

### Increased ICMS frequency leads to increased perceived magnitude

Three lines of evidence suggest that higher ICMS frequencies evoke more intense percepts. First, detection thresholds are lower indicating increased sensitivity at higher frequencies (Kim et al., 2015). Second, animals exhibit a highly consistent bias to select the higher-amplitude stimulus as being higher in frequency, even though this leads to less reward (Figure 3, Figure 4, Supplementary figure 10). This systematic bias implies a systematic relationship between frequency and perceived magnitude. Third, on experimental blocks comprising primarily equal-amplitude stimuli, catch trials – when both stimuli had the same frequency but different amplitudes – resulted in the systematic selection of the higher-amplitude stimulus, consistent with the hypothesis that the animal relied in part on intensity to make these discrimination judgments (Supplementary figure 10). Note that systematic selection biases on catch trials disappeared to the extent that animals judged frequency independently of amplitude (Figure 4, see below). Increased perceived magnitude at higher frequencies has also been reported for peripheral nerve stimulation (D’Anna et al., 2017; Graczyk et al., 2016) and is consistent with the hypothesis that intensity is determined by the spike rate evoked in the neuronal population (Kim et al., 2017).

### Changes in ICMS frequency can lead to changes in the quality of the evoked percept

On a number of electrodes, animals could select the higher frequency stimulus even when (1) it was much lower in amplitude and perceived as less intense than the stimulus with which it was paired and (2) it was interleaved with other pairs in which the higher frequency stimulus was perceived as more intense. In these cases, when the animal was presented with catch trials – in which both stimuli were of equal frequency but different amplitude – animals did not exhibit a bias to select the higher-amplitude stimulus (Figure 4, Supplementary figure 10). The animal thus demonstrated that it was largely ignoring differences in magnitude in making its frequency judgments. Furthermore, on those electrodes, the animal’s behavior was inconsistent with the hypothesis that frequency and amplitude affect a common sensory continuum – magnitude (Supplementary Figures 6 and 7). Indeed, the amplitude bias was weaker than would be predicted based on single-parameter discrimination performance (amplitude or frequency).

Together, these observations are consistent with the hypothesis that changes in frequency also affect the quality of the evoked percept on some electrodes. That the animals were often able to transfer performance from one electrode to the next suggests that the effect of frequency on sensory quality was consistent across electrodes. Indeed, had the perceptual effect been very different from electrode to electrode, the animal would have had to discover the relevant sensory continuum on an electrode by electrode basis. The inability to distinguish frequency independently of amplitude on other electrodes indicates that the effect of frequency at those electrodes is indistinguishable from that of amplitude, is subtle and drowned out by the fluctuations in amplitude, or does not lie on a discernible continuum.

### Neural codes

Any proposed neural code to explain the discrimination behavior must account for the animals’ ability to distinguish increases in frequency from increases in amplitude despite the fact that both stimulation parameters affect the firing rate of neural populations near the electrode tip. While higher amplitudes lead to the recruitment of a larger volume of neurons (Tehovnik, 1996; Tolias et al., 2005) and to changes in the spatial distribution of neuronal activity (Histed et al., 2009), the precise shape of recruitment is frequency dependent (Michelson et al., 2018). In principle, then, the spatial pattern of neuronal activation may have shaped the resulting percept and mediated the animals’ frequency discrimination behavior. We ruled out this possibility – and others positing that frequency discrimination relied on a spatial pattern of activation – by showing that performance is largely unhindered by random changes in amplitude from pulse to pulse, leading to a spatial pattern of activation that varies from pulse to pulse.

Having excluded the population spike rate and spatial hypotheses, we propose that frequency discrimination relies on the temporal structure of the ICMS-evoked activity. Indeed, each ICMS pulse synchronously activates a large population of neurons around the electrode tip (Tehovnik, 1996), so periodic stimulation results in synchronized periodic responses across a large swath of somatosensory cortex. The animals’ ability to discriminate ICMS frequency relies on the ability to detect differences in the temporal patterning in the population response, which in turn result in differences in sensation quality along a continuum that is distinct from magnitude. According to this hypothesis, the drop-off in performance for frequencies above 200Hz is caused by an inability of neuronal populations to phase lock at these frequencies. Note that a small population of neurons in somatosensory cortex has been shown to phase lock to vibratory stimuli up to 800 Hz (Harvey et al., 2013), but it is unlikely that hundreds or thousands of neurons could do so. According to this hypothesis, differences across electrodes would be due to differences in the ability of different local circuits to phase lock to stimulation, a hypothesis that can be tested by recording the electrically induced neuronal activity.

### Implications for neuroprosthetics

Regardless of the relevant perceptual continuum and neural mechanisms, ICMS frequency exerts a robust influence on the evoked percept. Indeed, we estimate that the perceptually relevant range of frequencies (from 10 to 200 Hz) accommodates 10 to 20 non-overlapping JNDs. That is, changes in frequency can lead to tens of mutually discriminable percepts. In contrast, amplitude JNDs – which range from 14 to 30 µA (Flesher et al., 2016; Kim et al., 2015) – provide at best 5 to 7 mutually discriminable percepts, from detection threshold (around 20-30 µA at 300 Hz) to the maximum amplitude used for human experiments (100 µA) (Flesher et al., 2016; Kim et al., 2015; Tabot et al., 2013).

To the extent that ICMS frequency and amplitude have different sensory correlates, the sensory space that can be achieved by changing these two stimulation parameters is vast (consisting of up to 140 discriminable percepts). Note, however, that frequency and amplitude can never be completely dissociated, even after extensive training on the best electrodes. More importantly, they cannot be dissociated at all on some electrodes. The challenge, then, will be to harness these two parameters in the most effective way, which will require taking into consideration the stark inter-electrode differences.

## Methods

### Animals

Three male Rhesus macaques (Macaca *mulatta*), ranging in age from 7 to 9 years old and weighing between 9 and 10 kg, participated in this study. Animal care and handling procedures were approved by the University of Chicago Animal Care and Use Committee.

### Implants

Each animal was implanted with one Utah electrode array (UEA, Blackrock Microsystems, Inc., Salt Lake City, UT) in the hand representation of area 1 (Figure 1). Each UEA consists of 96 1.5mm-long electrodes with tips coated in iridium oxide, spaced 400 µm apart, and spanning 4 mm × 4 mm of the cortical surface. The hand representation in area 1 was targeted based on anatomical landmarks. Given that the arrays were continuous to the central sulcus and area 1 spans approximately 3-5 mm of cortical surface from the sulcus (Pons et al., 1985), few if any electrodes were located in area 2. Given the length of the electrodes, their tips likely terminated in the infragranular layers of somatosensory cortex if embedded to their base, as we have previously shown in postmortem histological analysis with other animals instrumented with identical arrays (Rajan et al., 2015). We mapped the receptive field of each electrode by identifying which areas of skin evoked significant z-scored multiunit activity (Callier et al., 2018).

### Stimuli

Intracortical stimulation (ICMS) consisted of cathodal phase-leading symmetrical biphasic pulses delivered through a 96-channel neurostimulator (CereStim R96, Blackrock Microsystems Inc., Salt Lake City, UT). Across all tested stimulation regimes, pulse train frequencies ranged 17 to 400 Hz, pulse amplitudes ranged from 44 to 100 µA, and phase durations equaled 200 or 400 µs. The interval between phases was always 53 µs. All biphasic pulses within the same stimulus were separated by the same time interval (pulse trains were periodic). In some experiments, all the pulses in a train had the same amplitude, in others, amplitude varied from pulse to pulse (see below).

### Behavioral task

The animals were seated at the experimental table facing a monitor, which signaled the trial progression (Figure 1). Eye movements were tracked with an optical eye-tracking system (MR PC60, Arrington Research, Scottsdale, AZ). The animals initiated trials by directing their gaze to a cross in the center of the monitor. A trial was aborted if an animal failed to maintain its gaze on the center until the appearance of response targets. Each trial comprised two successive stimulus intervals, each indicated by a circle on the video monitor, lasting 1 s, and separated by a 1-s interstimulus interval during which the circle disappeared, followed by a response interval during which two response targets appeared on either side of the gaze fixation point (Figure 1). The animals’ task was to judge which of the two pulse trains was higher in frequency. The animals responded by making saccadic eye movements toward the left (selecting the first stimulus) or right (second stimulus) target. Correct responses were rewarded with a drop of juice. Psychophysical performance was calculated as the proportion of trials on which the higher frequency stimulus was selected.

### Experimental design

#### Stimulus set for detailed psychometric curves

we first performed extensive testing on a small group of electrodes, building psychometric curves spanning a wide range of frequencies. In each test block, consisting of several hundred trials, one stimulus of each pair had the same standard frequency (20, 50, 100, or 200 Hz) and the other stimulus was drawn from a set of comparison frequencies around the standard (comparisons for 20Hz standard: 17, 18, 22, 23, 26, 29 Hz; comparisons for 50Hz standard: 30, 35, 40, 45, 55, 60, 65, 70 Hz; comparisons for 100Hz standard: 50, 75, 150, 200, 250, 300 Hz; comparisons for 200Hz standard: 50, 100, 150, 250, 300, 350, 400 Hz). Phase duration during each pulse was 200 µs in the higher frequency range (100 and 200 Hz standards) and 400 µs in the lower frequency range (20 and 50 Hz standards) to ensure that the stimuli were well above detection level *(1)*. Stimulus amplitudes were 50, 60, 70, or 80 µA. Every possible combination of frequencies (standard vs. comparisons) and amplitudes, numbering hundreds of unique stimulus pairs, was presented in each test block, ensuring the animals had to perform frequency discrimination instead of memorizing the correct responses to individual pairs of stimuli. For each electrode/standard combination, animals were trained until they reached stable performance, a process which could take weeks or even months as we incrementally included harder stimulus pairs (those with small frequency differences and large amplitude confounds). We over-represented stimulus pairs in which the higher frequency stimulus had a lower amplitude to discourage the animals’ reliance on perceptual magnitude in making their frequency judgments. The psychometric curves were constructed after asymptotic performance was achieved. We extensively tested 5 electrodes (4 from monkey A, 1 from monkey B) at standard frequencies of 50, 100, and 200Hz. Only 1 electrode from each monkey was tested at the 20 Hz standard before testing was cut short by the failure of Monkey A’s array. 3 electrodes from monkey C were extensively tested at only the 100Hz standard before testing was cut short by health issues which precluded water restriction.

#### Reduced stimulus set

After extensively testing a few electrodes, which took weeks or even months for each electrode and standard frequency, we developed a reduced stimulus subset to test the animals’ performance at a faster pace over a wide range of electrodes. In this stimulus set, the frequency difference was always 100 Hz, a salient difference according to the psychometric curves obtained from the full set, and the base (lower) frequency was 70, 80, 90, 100, 110, 120, 130, or 170 Hz. The amplitudes were the same as in the full set (50, 60, 70, or 80 µA) and, again, parametrically combined. Here too, we over-represented stimulus pairs in which the higher frequency stimulus had a lower amplitude to reduce the animals’ reliance on intensive cues in making their frequency judgments. We tested 17 electrodes from monkey B with this stimulus set. Instead of first training to asymptotic performance while incrementally adding harder stimulus pairs, we had the animal complete several thousand trials (from 2500 to 6000) with the complete stimulus set at each electrode. On 4 of the 17 electrodes, the animal performed the frequency discrimination task on 2 subsets of stimuli in separate experimental blocks: one containing only pairs with equal amplitudes and one containing only pairs in which the base stimulus’ amplitude was 30 µA higher than that of the comparison stimulus (Supplementary figure 10).

#### Catch trials

Included in the reduced stimulus set was a small proportion of trials (∼5%) on which the two stimuli in the pair were at the same frequencies but differed in amplitude by 30 µA. On these catch trials, all base frequencies were used. The animal was rewarded randomly during these trials. The animal’s bias toward the higher or lower amplitude stimulus in the absence of frequency differences gauged its reliance on intensive cues (Figure 4, Supplementary figure 10).

#### Variable amplitude pulse trains

The mean amplitude of the pulse train was of 58, 72, or 86µA but the amplitudes of individual pulses spanned a range around the mean (44 to 72 µA, 58 to 86 µA, or 72 to 100 µA in 2-µA increments), presented in random order. One of the stimuli in each pair as at 75, 105, 135, or 165 Hz and the other was 90 Hz higher. For each pair of frequencies, every possible combination of amplitudes (variable or constant) was tested. Of the 17 electrodes tested with the reduced stimulus set, 4 showing small effects of amplitude on performance and 3 showing large amplitude effects were tested with variable amplitude pulse trains.

#### Detection threshold amplitudes

One interval on each trial contained a 100-Hz pulse train at 10, 25, 40, 55, or 70 µA and the other interval was empty. The animal reported which interval contained the stimulus. 12 electrodes from monkey B covering the range of susceptibilities to amplitude were tested.

### Data analysis

#### Psychophysics

We built psychometric curves by fitting performance at each comparison frequency to a cumulative normal density function. Just-noticeable-differences (JNDs) and Weber fractions were calculated using only trials on which both the stimuli in the pair were equal in amplitude using a criterion performance of 75% correct (Figure 2). JNDs were calculated as the average of the frequency differences required for threshold performance above and below the standard frequency. These were nearly equal for standard frequencies of 20 and 50 Hz but tended to be asymmetric at higher frequencies (with the upper JND greater than the lower one). If performance did not reach threshold for comparison frequencies above the standard for a given electrode and standard, only the frequency difference below the standard was used. This only occurred with the 200-Hz standard frequency. When considering the effect of amplitude differences on the animals’ choices, points of subjective equality (PSEs) were computed for each amplitude difference between comparison and standard stimuli (ranging from −30µA to 30µA). The PSE is the comparison frequencies at which the probability of the animal selecting the comparison was 0.5. As PSE exhibited a relationship with amplitude difference that was well approximated by a logarithmic function, we fit a line through the natural log of the PSEs at each amplitude difference to obtain a slope which quantified the equivalence tradeoff between frequency and amplitude. A small slope would result from PSEs that were near each other, and would indicate that amplitude had a relatively small effect on the animal’s choices because only a small change in frequency was required to offset a 1µA difference in amplitude. We computed these slopes for every electrode at every standard frequency.

#### Gauging the relative contribution of amplitude and frequency to discrimination judgments

The behavioral data show that the animals’ judgments depended on both frequency and amplitude, but the relative contribution of these two stimulation parameters varied from electrode to electrode. To assess the relative contributions of frequency and amplitude to discrimination judgments, we modeled the position of each stimulus along the task-relevant sensory dimension as a weighted combination of the stimulus’ frequency and amplitude. To predict the performance of the animal, we first subtracted the value of the standard from that of the comparison along this sensory continuum for each pair. The resulting differences were then the input to a sigmoid (cumulative normal density function). The resulting function comprised 3 free parameters: two regression weights (frequency, amplitude) and one sigmoid parameters (standard deviation). For each of the five electrode through which the complete stimulus set was delivered, the function was optimized to predict the animal’s behavioral performance. The model provided an accurate fit of the behavioral data (*R*^*2*^ = mean ± s.e.m., Supplementary figure 6B). The regression weights gauge the relative contribute of frequency and amplitude in determining the animals’ choices (Supplementary figure 6B). For example, to the extent that the regression weight for amplitude was low, we concluded that the animal was able to discriminate frequency independent of amplitude on that electrode.

#### Generating equivalent frequency-amplitude tradeoffs using single-variable discrimination

We wished to test the hypothesis that frequency discrimination performance was based entirely on intensive cues. That is, changes in frequency and changes in amplitude have the same effect on the evoked percept. To this hypothesis, we first assessed discriminability when only frequency or only amplitude changed and assessed whether the performance in these experiments can account for performance when both parameters varied (Supplementary figure 7, Supplementary figure 8). To gauge sensitivity to ICMS amplitude, we used previously published data (Kim et al., 2015). In these experiments, the standard amplitude was 70µA, and the comparisons were 40, 50, 60, 80, 90, and 100 µA. Psychometric curves were built based on amplitude discrimination performance averaged across all frequencies (50, 100, 250, and 500 Hz) because we found that frequency had a negligible effect on amplitude JNDs (Kim et al., 2015). To gauge sensitivity to ICMS frequency, we restricted the analysis to trials in which both stimuli had amplitudes of 70 µA.

Using the psychometric functions derived from the single-parameter discrimination experiments, we computed equivalent frequency-amplitude tradeoffs by equating changes in frequency and amplitude that resulted in equal discrimination performance. For example if an amplitude difference of ±10 µA and a frequency difference of ±8 Hz both resulted in a discrimination performance of 65%, ±10 µA and ±8 Hz were considered to be perceptually equivalent. The resulting predicted equal intensity curves were smoothed the tangent was computed at each standard frequency and at 70µA. The frequency/amplitude tradeoff implied by this slope could then be compared to the tradeoff obtained with the linear model above, which was derived without the assumption that the sensory consequences of amplitude and frequency changes are indistinguishable. Because the amplitude discrimination experiment was performed on different electrodes (8 total, collectively the “amplitude electrodes”) than the frequency experiment (the “frequency electrodes”), we matched each “frequency electrode” with each of the 8 “amplitude electrodes” in turn and carried out this analysis. This yielded a distribution of possible tangent slopes based on each “frequency electrode’s” sensitivity to amplitude. If the frequency-amplitude tradeoff computed from the model fell outside of this distribution, the results from this analysis were inconsistent with the hypothesis that frequency and amplitude only affect a common intensive continuum.

## Acknowledgments

This work was supported by NINDS grant NS095251.

## Supplementary text

### Validation of catch trials

On four electrodes, we trained monkey B to perform frequency discrimination on 2 subsets of stimuli before using the full stimulus set: one with both stimuli in the pair of equal amplitude and one in which the higher frequency stimulus always was at a much lower amplitude. High performance was achieved on all four electrodes for these two subsets (Supplementary figure 10). However, catch trials revealed that the animal was biased toward more intense stimulus in the first set but toward less intense stimuli in the second set. These results are consistent with the hypothesis that catch trials reveal the animal’s reliance on intensive cues and indicate that the 30-µA amplitude difference was sufficient to overcome the 100-Hz frequency difference such that the higher frequency stimulus felt less intense than the lower frequency one. When the animal was faced with the full set (which comprised stimuli from the first two), the animal was able to perform the task correctly regardless of amplitude differences on only two of the four electrodes (Supplementary figure 10).

### Learning during task transfer

All the data described above reflect monkey B’s asymptotic performance, sometime after weeks of training on each electrode and standard frequency. Next, we examined how the animal’s performance evolved during training. We found that, on some electrodes, the animal appeared unable to overcome the amplitude confound over the course of up to 6000 trials (example 2 in Supplementary figure 11). On other electrodes, the animal relied on intensity cues at first, but learned over an extended period to discriminate frequency independent of amplitude (example 1). On yet other electrodes, the animal’s performance was initially high but somewhat amplitude-dependent and became, relatively rapidly, independent of amplitude (example 3). We found that the majority of electrodes yielded good performance immediately (above criterion of 75% at all amplitude differences) or never yielded good performance (the animal did not learn to perform above criterion at all amplitude differences within 6000 trials)(Supplementary figure 11B).

### Adaptation index

We developed an adaptation Index to quantify the relative contributions of slowly adapting and rapidly adapting signals to the multi-unit responses to skin indentations at each electrode (Supplementary figure 4), as has been previously done (Callier et al., 2018; Pei et al., 2009). For this calculation, we normalized neural responses during sustained indentation and indentation offset to their respective grand means across electrodes. The adaptation index was then computed by dividing the normalized offset response by the sum of the normalized sustained and normalized offset responses for each electrode separately. An index of 0 denoted a pure SA1-like response, an index of 1 a pure RA-like response.

## Supplementary figures

**Supplementary figure 1:**
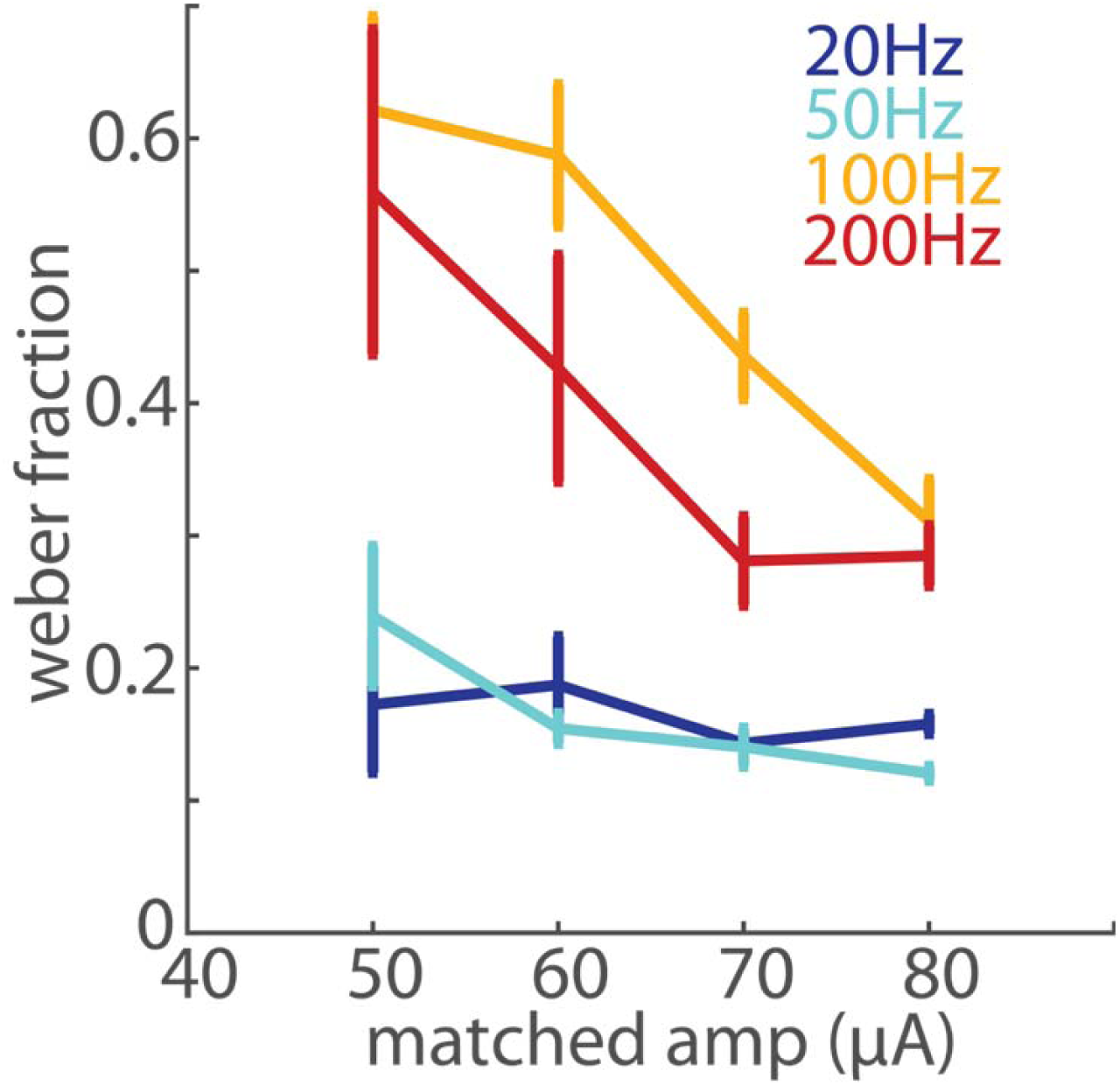
Weber fraction as a function of amplitude. Performance at each standard frequency is shown in a different color. Error bars show the standard error of the mean across the extensively tested electrodes from the three monkeys (2 electrodes from monkeys A and B for the 20Hz standard, 5 from monkeys A and B for the 50 and 200Hz standards, 8 electrodes the three monkeys for the 100Hz standard). Weber fractions were independent of amplitude in the lower frequency range, but decreased with amplitude for frequencies above 100 Hz.

**Supplementary figure 2:**
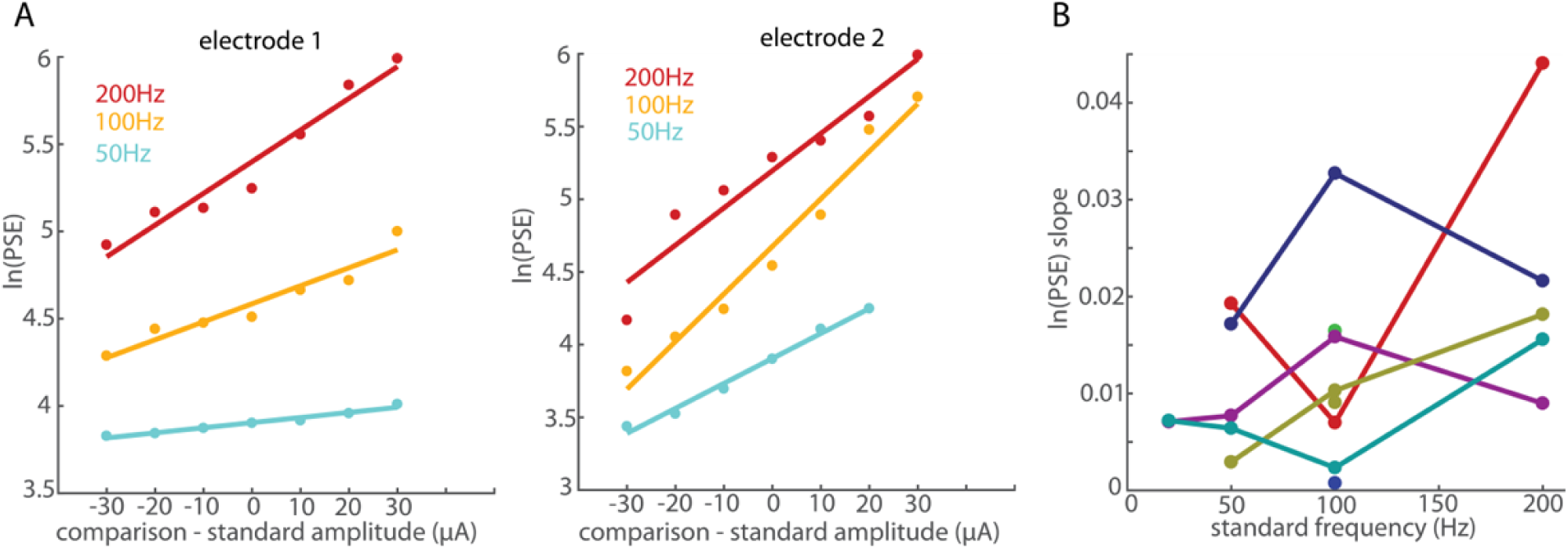
Quantifying the amplitude bias. A| Left: For the electrode shown in the top row of Figure 3, ln(PSE) vs. amplitude difference between the standard and comparison stimuli. Colors denote performance at different standard frequencies. The lines show the linear fit of ln(PSE) to the amplitude difference. Right: ln(PSE) for the electrode shown in the bottom row of Figure 3. The larger biasing effect of amplitude at the 50 and 100 Hz standards is reflected in the steeper slopes. B | Slopes vs. standard frequency for the 8 extensively tested electrodes. Different colors denote different electrodes.

**Supplementary figure 3:**
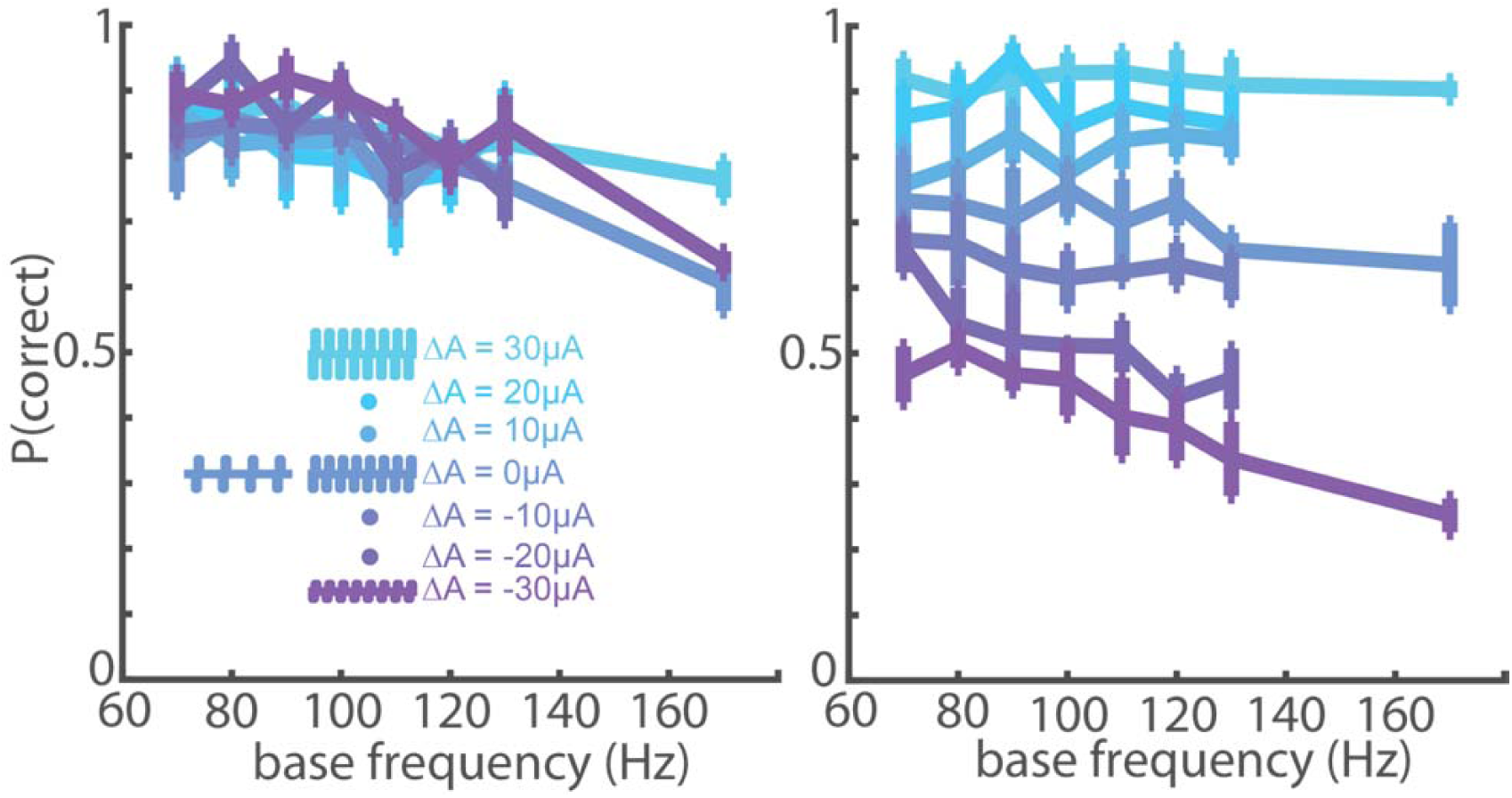
Performance as a function of base frequency for the 4 electrodes with the lowest spread (left 4 electrodes in figure 4) and the 4 electrodes with the greatest spread (right 4 electrodes in figure 4, excluding any marked with the green dot). Performance decreased slightly as the base frequency became larger.

**Supplementary figure 4:**
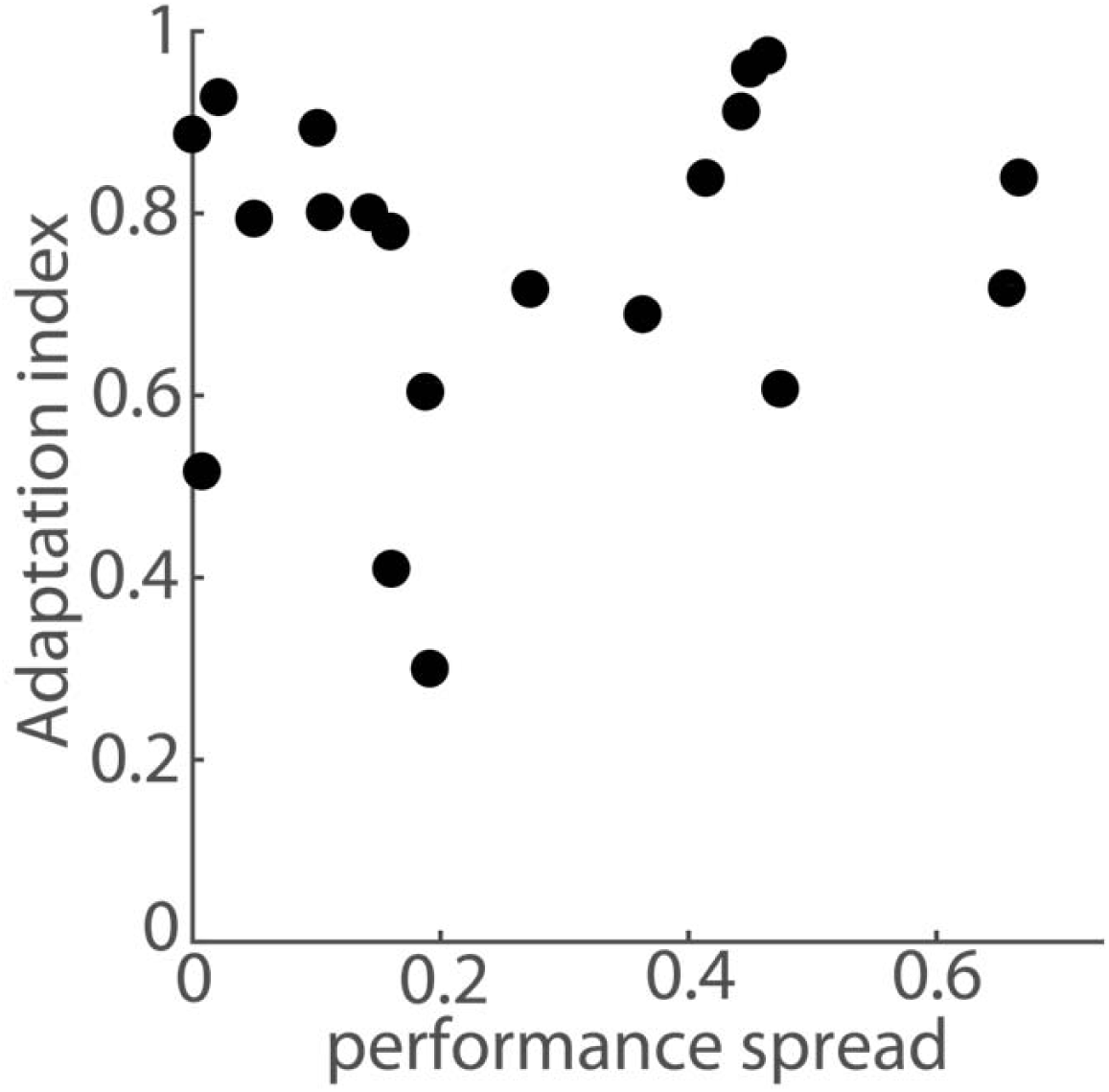
adaptation index as a function of the spread between performance at the two amplitude difference extremes (purple and cyan in figure 4). A lower spread indicates a greater ability to distinguish the effects of frequency and amplitude. There is no apparent relationship between performance at an electrode and the adaptation properties of the corresponding neural response (how “RA-like” it is). Adaptation index values could only be computed for 20 of the 25 electrodes shown in figure 4. No change in neural activity in response to indentations was recorded at the other 5 electrodes.

**Supplementary figure 5:**
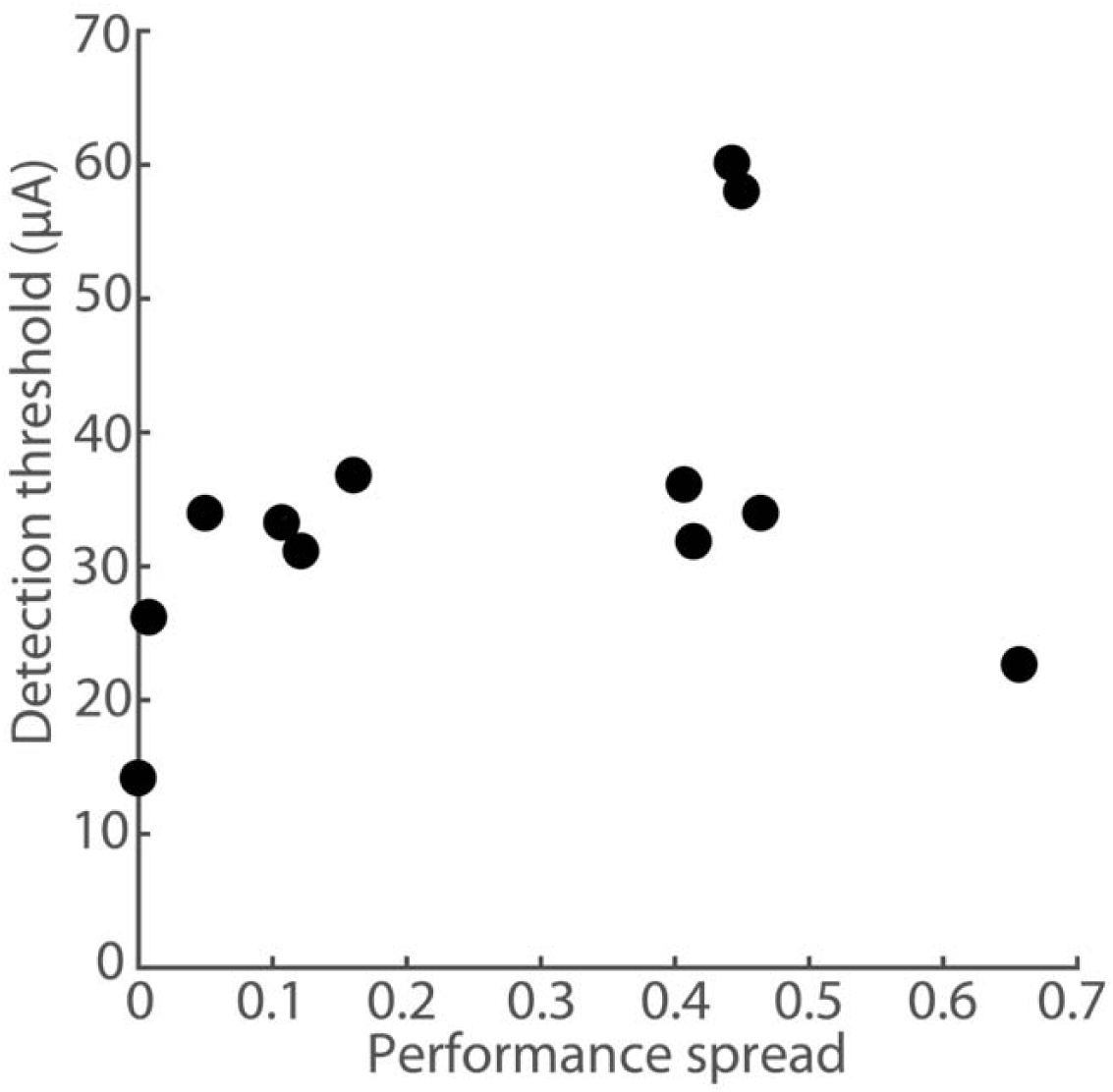
Detection threshold vs. susceptibility to the amplitude confound in the frequency discrimination task. Microstimulation frequency was 100Hz. The threshold is the minimum amplitude at which the animal detects the stimulus 75% of the time.

**Supplementary figure 6:**
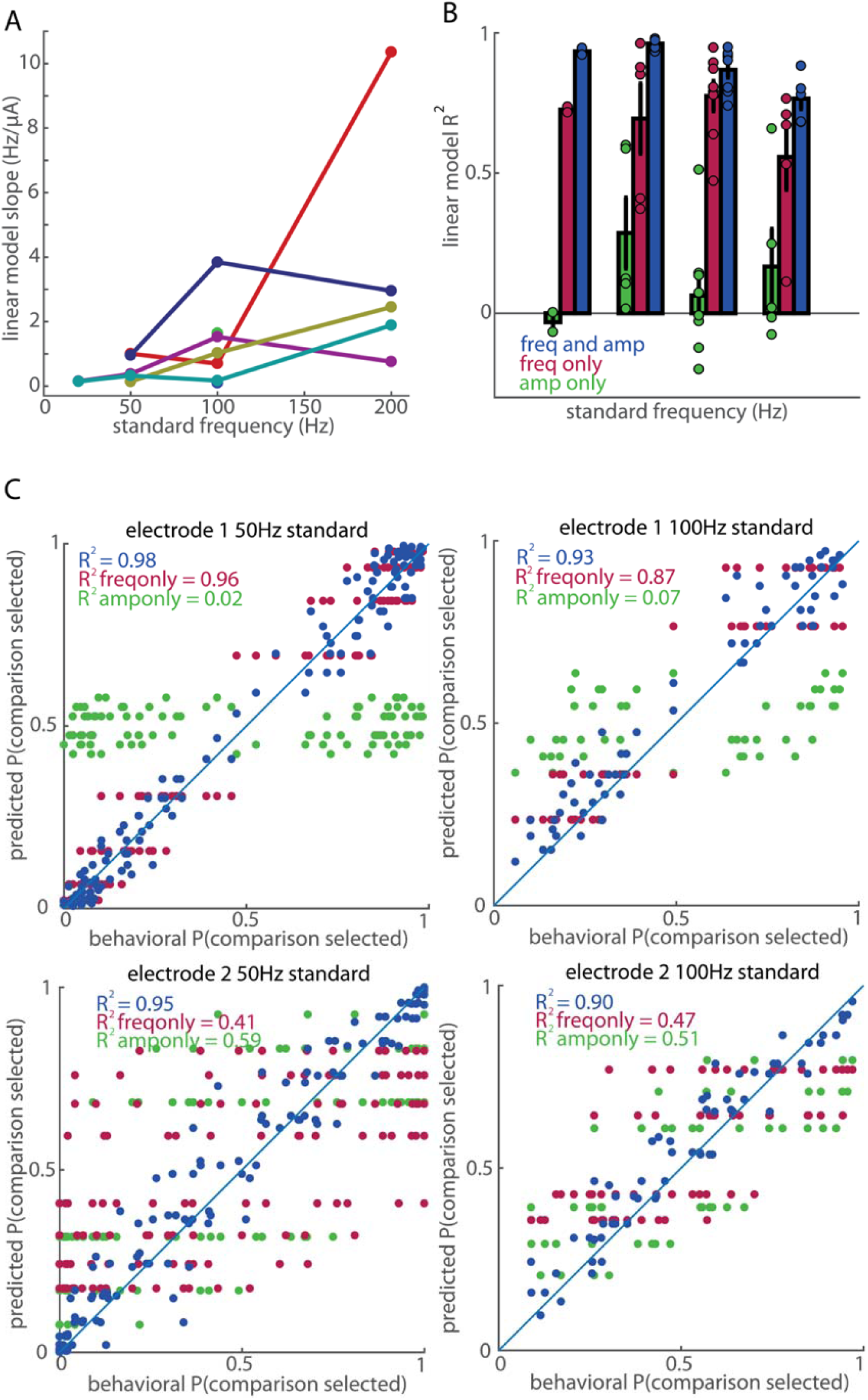
Contribution of frequency and amplitude to frequency discrimination performance. A| Slopes of equivalent frequency-amplitude tradeoffs (the number of Hz equivalent to a 1µA change), derived from the model fit, for each standard frequency. Different colors denote different electrodes. B| Goodness of fit of the model (*R*^*2*^) when using amplitude only (green), frequency only (red), or both (blue) as predictors. Bars denote the mean *R*^*2*^, error bars the standard error of the mean and dots the value for individual electrodes. C| For the same two electrodes and standard frequencies shown in figure 3, model predictions for each unique stimulus pair vs. actual performance. Amplitude dominated the animal’s choices on the bottom electrode to the point that the reconstruction using amplitude only is superior to the reconstruction using frequency only.

**Supplementary figure 7.**
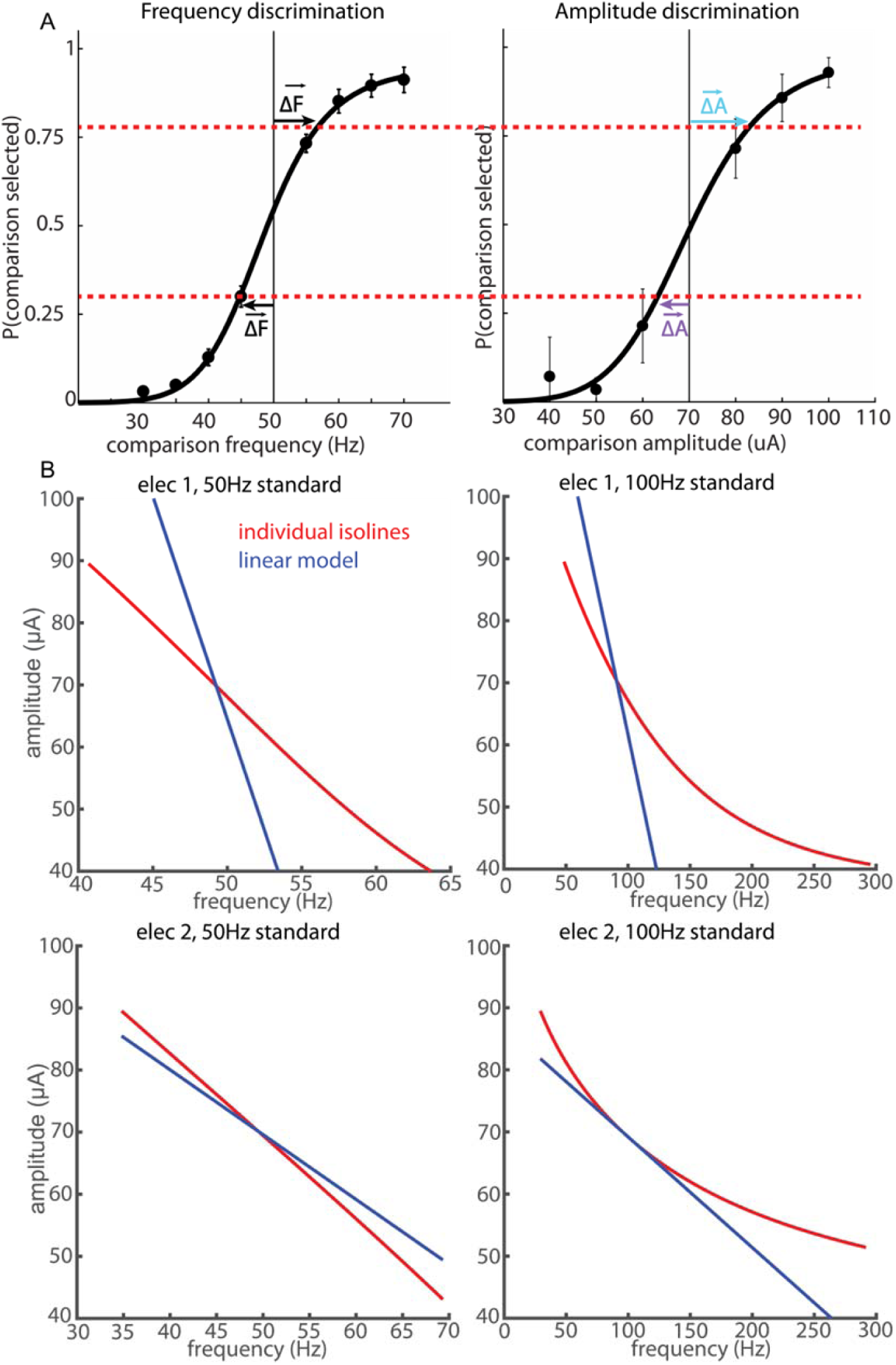
A| Construction of iso-intensity lines from single-variable discrimination experiments. Frequency differences in a frequency discrimination task (left) and amplitude differences in an amplitude discrimination task (right) that yield the same discrimination performance are used to obtain perceptual equivalence (red, below). Note that this methodology assumes that differences in amplitude and frequency affect the percept along the same sensory continuum. B| For the two electrodes and standards in figure 3, equivalence lines derived from single-variable discrimination (red) and from fitting behavior with the full stimulus set to a linear model (blue). At the standard frequency, the predicted equivalence line has a different slope than observed one for electrode 1, but not for electrode 2. The relative importance of frequency is much greater for electrode 1 than what could be expected from the single-variable experiment, indicating that amplitude and frequency affect the elicited sensation along different perceptual axes.

**Supplementary figure 8:**
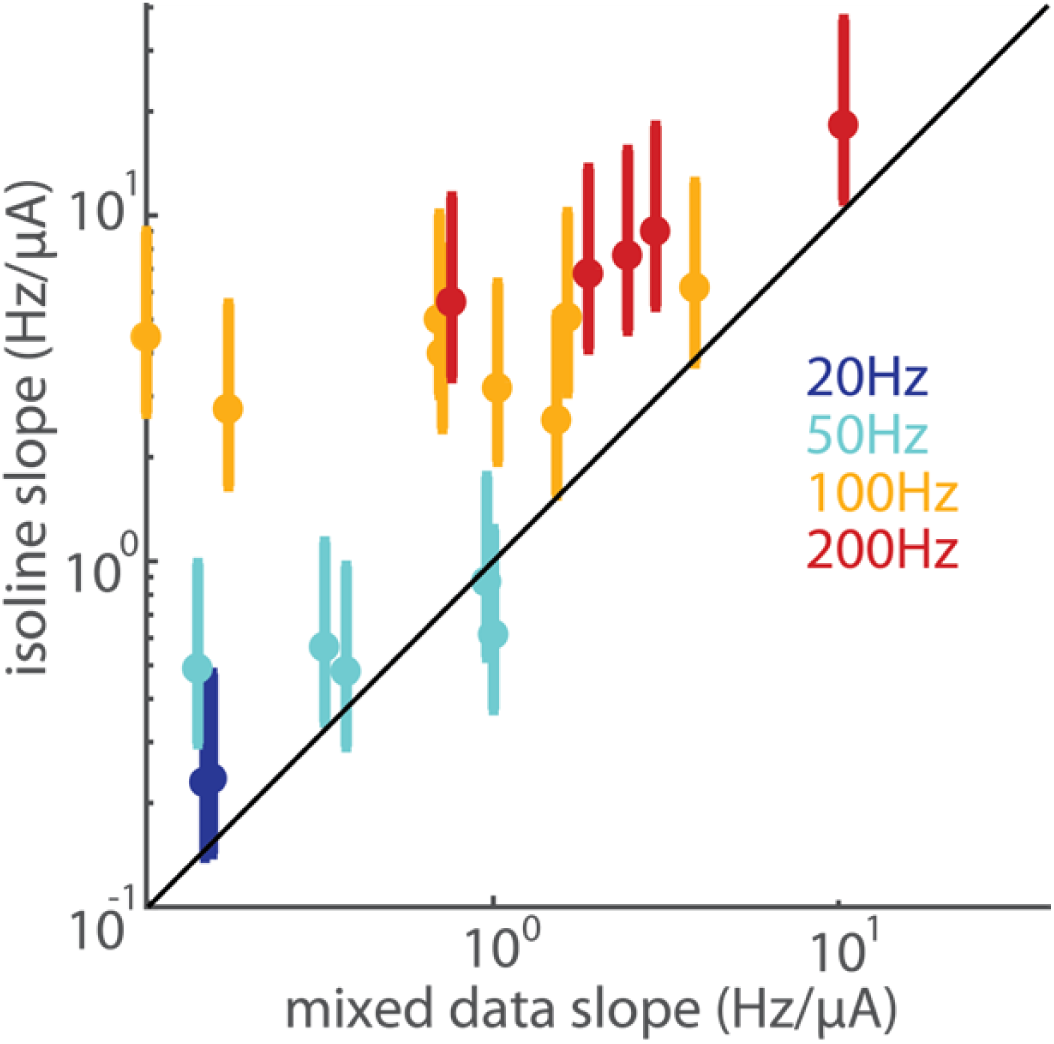
Equivalence slopes obtained using single-variable experiments vs equivalence slopes obtained from fitting a linear model to the frequency discrimination performance with variable amplitude, for the extensively studied electrodes at each standard frequency (2 for the 20Hz standard, 5 for the 50 and 200Hz standard, 8 for the 100Hz standard). The y axis error bars show the range of slopes obtained by matching each electrode’s frequency discrimination performance to the amplitude discrimination performances from many electrodes and the dot shows the slope obtained when the amplitude discrimination performances from the different electrodes are averaged together. The relative effect of frequency obtained from the mixed-variable experiment appears consistently greater, aside from two electrodes at the 50 Hz standard.

**Supplementary figure 9:**
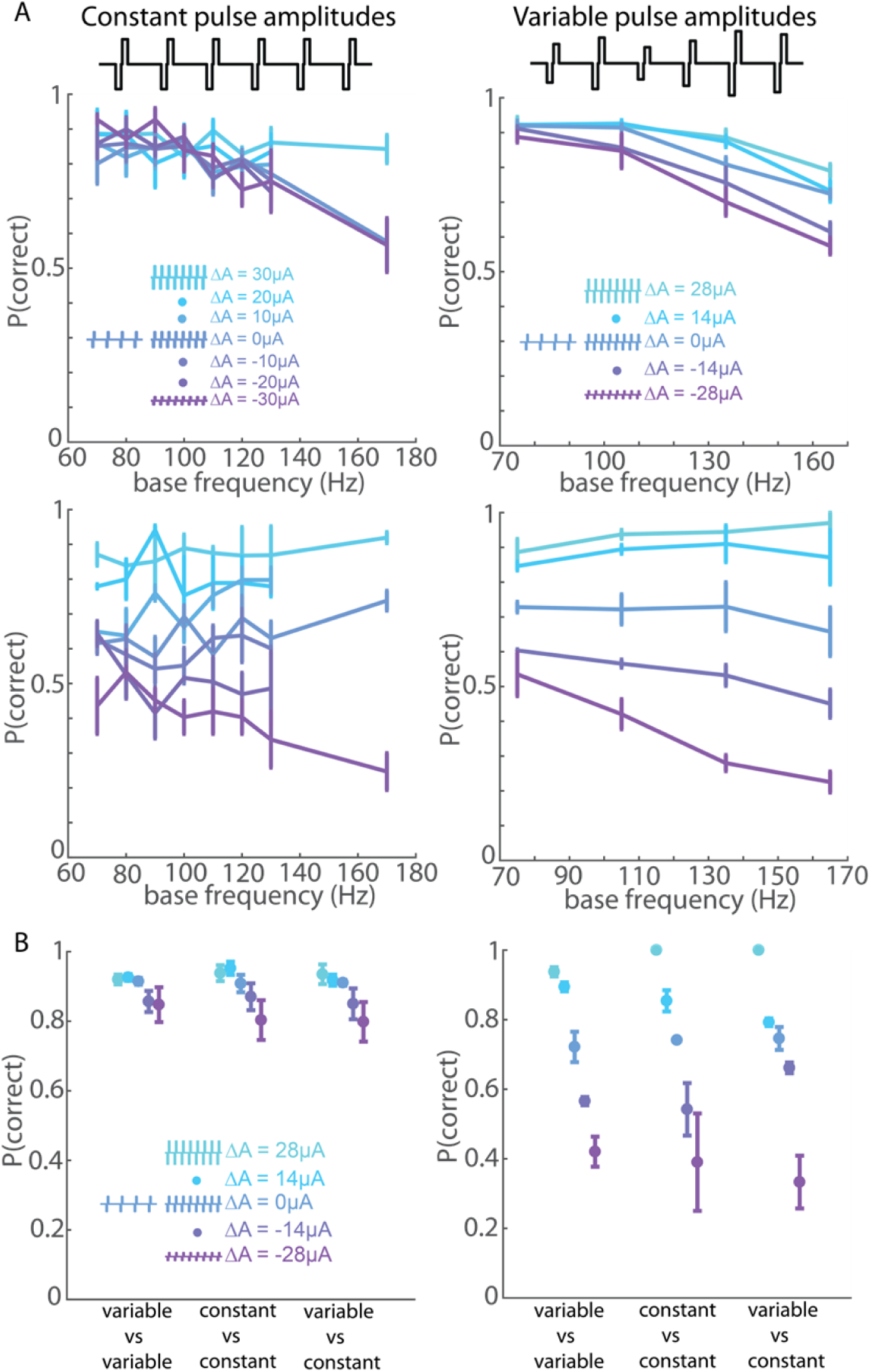
Varying individual pulse amplitude has a negligible effect on frequency discrimination performance. A| Monkey B’s performance vs. base frequency in the constant-amplitude experiment (left column) and the variable-amplitude experiment (right column) for the same 4 electrodes with weak amplitude bias (top row) and the same 2 electrodes with strong amplitude bias (bottom row). The frequency difference was always 100 Hz for the constant-amplitude experiment and 90 Hz for the variable-amplitude experiment due to hardware constraints (see methods). B| Performance when stimulus pulse trains were both variable-amplitude, were split, or were both constant-amplitude, for the 4 electrodes with weak amplitude bias (left) and the two with strong amplitude bias (right). Changing spatial distribution of the ICMS-induced activity on a pulse by pulse basis had little to no effect on the animal’s ability to discriminate frequency. The top right panel in A and the left panel in B are taken from figure 5.

**Supplementary figure 10.**
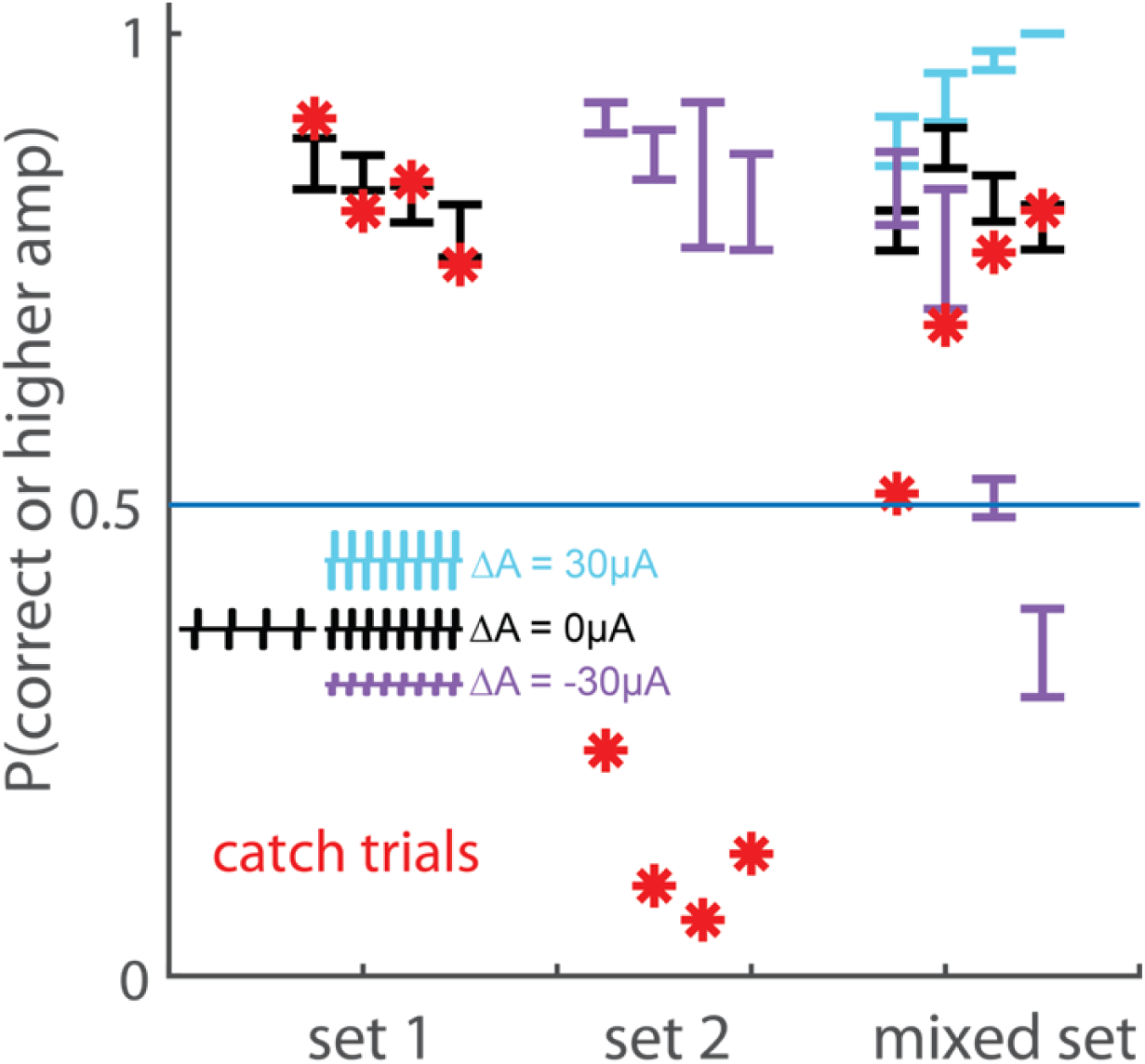
For four electrodes, p(correct) in the frequency discrimination task when amplitudes are matched (subset 1, black), when the higher frequency always has a lower amplitude (subset 2, purple), or when the full set is used (subset 3, only −30, 0, and 30 µA differences are shown). Red asterisks denote the probability of selecting the higher amplitude on catch trials.

**Supplementary figure 11:**
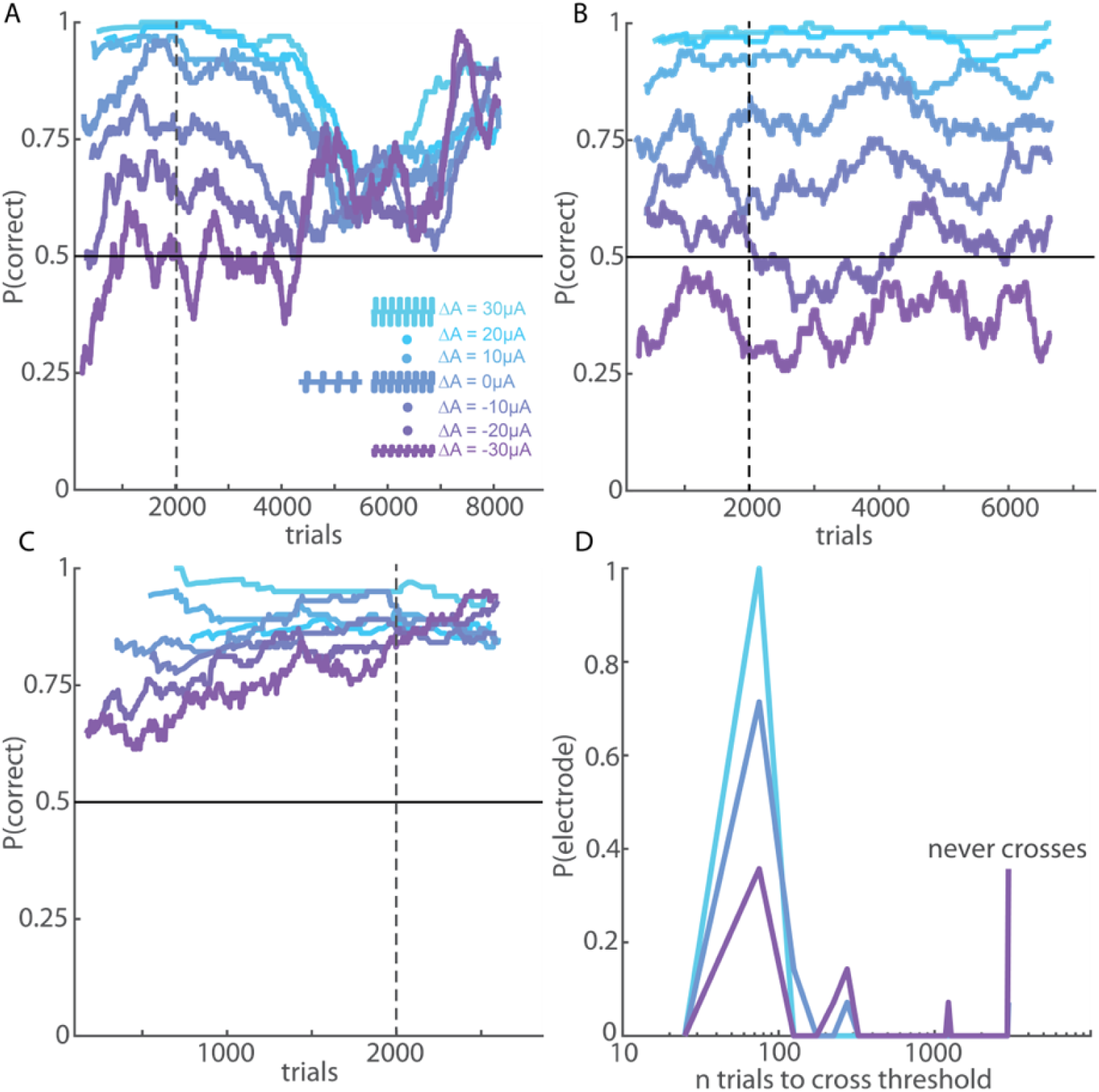
learning. A| Performance over time for an electrode on which the monkey first relied on intensity before learning to judge frequency independently. B| example of an electrode at which the animal continued to rely on intensity. C | example of an electrode at which the animal immediately performed well at all amplitude differences. D | Probability histogram of the number of trials of each condition required to achieve performance above a 75% threshold at amplitude differences of −30, 0, and 30µA (17 electrodes tested with the reduced stimulus set). When amplitudes were matched or the higher frequency had a higher amplitude, performance was above threshold immediately for all electrodes. When the lower frequency had a higher amplitude, at the vast majority of electrodes performance was either above threshold from the beginning, or never reached threshold. There were only a few electrodes for which performance in this condition was initially low but eventually met criterion performance.

## References

Bensmaia, S. J. (2015). Biological and bionic hands: Natural neural coding and artificial perception. Philosophical Transactions of the Royal Society B: Biological Sciences, 370(1677). https://doi.org/10.1098/rstb.2014.0209

Berg, J. A., Dammann, J. F., Tenore, F. V., Tabot, G. A., Boback, J. L., Manfredi, L. R., … Bensmaia, S. J. (2013). Behavioral demonstration of a somatosensory neuroprosthesis. IEEE Transactions on Neural Systems and Rehabilitation Engineering, 21(3), 500–507. https://doi.org/10.1109/TNSRE.2013.2244616

Callier, T., Suresh, A. K., & Bensmaia, S. J. (2018). Neural Coding of Contact Events in Somatosensory Cortex. Cerebral Cortex, 1–15. https://doi.org/10.1093/cercor/bhy337

D’Anna, E., Petrini, F. M., Artoni, F., Popovic, I., Simanić, I., Raspopovic, S., & Micera, S. (2017). A somatotopic bidirectional hand prosthesis with transcutaneous electrical nerve stimulation based sensory feedback. Scientific Reports, 7(1), 1–15. https://doi.org/10.1038/s41598-017-11306-w

Flesher, S. N., Collinger, J. L., Foldes, S. T., Weiss, J. M., Downey, J. E., Tyler-Kabara, E. C., … Gaunt, R. A. (2016). intracortical microstimulation of human somatosensory cortex. Sci. Transl. Med, 8, 361–141. https://doi.org/10.1126/scitranslmed.aaf8083

Graczyk, E. L., Schiefer, M. A., Saal, H. P., Delhaye, B. P., Bensmaia, S. J., & Tyler, D. J. (2016). The neural basis of perceived intensity in natural and artificial touch. Science Translational Medicine, 8(362), 1–11. https://doi.org/10.1126/scitranslmed.aaf5187

Harvey, M. A., Saal, H. P., Dammann, J. F., & Bensmaia, S. J. (2013). Multiplexing Stimulus Information through Rate and Temporal Codes in Primate Somatosensory Cortex. PLoS Biology, 11(5). https://doi.org/10.1371/journal.pbio.1001558

Histed, M. H., Bonin, V., & Reid, R. C. (2009). Direct Activation of Sparse, Distributed Populations of Cortical Neurons by Electrical Microstimulation. Neuron, 63(4), 508–522. https://doi.org/10.1016/j.neuron.2009.07.016

Kim, S., Callier, T., & Bensmaia, S. J. (2017). A computational model that predicts behavioral sensitivity to intracortical microstimulation. Journal of Neural Engineering, 14(1). https://doi.org/10.1088/1741-2552/14/1/016012

Kim, S., Callier, T., Tabot, G. A., Gaunt, R. A., Tenore, F. V., & Bensmaia, S. J. (2015). Behavioral assessment of sensitivity to intracortical microstimulation of primate somatosensory cortex. Proceedings of the National Academy of Sciences of the United States of America, 112(49). https://doi.org/10.1073/pnas.1509265112

Kim, S., Callier, T., Tabot, G. A., Tenore, F. V., & Bensmaia, S. J. (2015). Sensitivity to microstimulation of somatosensory cortex distributed over multiple electrodes. Frontiers in Systems Neuroscience, 9(APR). https://doi.org/10.3389/fnsys.2015.00047

Michelson, N. J., Eles, J. R., Vazquez, A. L., Ludwig, K. A., & Kozai, T. D. Y. (2018). Calcium activation of cortical neurons by continuous electrical stimulation: Frequency dependence, temporal fidelity, and activation density. Journal of Neuroscience Research, (September 2018), 620–638. https://doi.org/10.1002/jnr.24370

Pei, Y.-C., Denchev, P. V., Hsiao, S. S., Craig, J. C., & Bensmaia, S. J. (2009). Convergence of Submodality-Specific Input Onto Neurons in Primary Somatosensory Cortex. Journal of Neurophysiology, 102(3), 1843–1853. https://doi.org/10.1152/jn.00235.2009

Pons, T. P., Garraghty, P. E., Cusick, C. G., & Kaas, J. H. (1985). A sequential representation of the occiput, arm, forearm and hand across the rostrocaudal dimension of areas 1, 2 and 5 in macaque monkeys. Brain Research, 335(2), 350–353. https://doi.org/10.1016/0006-8993(85)90492-5

Rajan, A. T., Boback, J. L., Dammann, J. F., Tenore, F. V., Wester, B. A., Otto, K. J., … Bensmaia, S. J. (2015). The effects of chronic intracortical microstimulation on neural tissue and fine motor behavior. Journal of Neural Engineering, 12(6). https://doi.org/10.1088/1741-2560/12/6/066018

Romo, R., Hernandez, A., & Zainos, A. (2000). Sensing without Touching : Somatosensory discrimination based on cortical microstimulation. Neuron, 26, 273–278.

Romo, R., Hernández, A., Zainos, A., & Salinas, E. (1998). Somatosensory discrimination based on cortical microstimulation. Nature, 392(6674), 387–390. https://doi.org/10.1038/32891

Saal, H. P., & Bensmaia, S. J. (2014). Touch is a team effort: Interplay of submodalities in cutaneous sensibility. Trends in Neurosciences, 37(12), 689–697. https://doi.org/10.1016/j.tins.2014.08.012

Salas, M. A., Bashford, L., Kellis, S., Jafari, M., Jo, H., Kramer, D., … Andersen, R. A. (2018). Proprioceptive and cutaneous sensations in humans elicited by intracortical microstimulation. ELife, 7, 1–11. https://doi.org/10.7554/eLife.32904

Tabot, G. A., Dammann, J. F., Berg, J. A., Tenore, F. V., Boback, J. L., Vogelstein, R. J., & Bensmaia, S. J. (2013). Restoring the sense of touch with a prosthetic hand through a brain interface. Proceedings of the National Academy of Sciences, 111(2), 875–875. https://doi.org/10.1073/pnas.1322627111

Tehovnik, E. J. (1996). Electrical stimulation of neural tissue to evoke behavioral responses. Journal of Neuroscience Methods, 65 VN-r(1), 1–17. https://doi.org/10.1016/0165-0270(95)00131-X

Tolias, A. S., Sultan, F., Augath, M., Oeltermann, A., Tehovnik, E. J., Schiller, P. H., & Logothetis, N. K. (2005). Mapping cortical activity elicited with electrical microstimulation using fMRI in the macaque. Neuron, 48(6), 901–911. https://doi.org/10.1016/j.neuron.2005.11.034

